# Experimental control of macrophage pro-inflammatory dynamics using predictive models

**DOI:** 10.1101/826966

**Authors:** Laura D. Weinstock, James E. Forsmo, Alexis Wilkinson, Jun Ueda, Levi B. Wood

**Affiliations:** Parker H. Petit Institute for Bioengineering & Bioscience, Georgia Institute of Technology, Atlanta, GA 30332; The Wallace H. Coulter Department of Biomedical Engineering, Georgia Institute of Technology, Atlanta, GA 30332; School of Chemical & Biomolecular Engineering, Georgia Institute of Technology, Atlanta, GA 30332; George W. Woodruff School of Mechanical Engineering, Georgia Institute of Technology, Atlanta, GA 30332

## Abstract

In numerous diseases, dysregulated macrophage polarization can not only impair recovery, but can also promote further injury and pathogenesis. Thus, a method of predicting and dynamically controlling macrophage polarization will enable a new strategy for treating diverse inflammatory diseases. Here, we developed a model-predictive control framework to temporally regulate macrophage polarization. Using RAW 264.7 macrophages as a model system, we enabled temporal control by identifying transfer function models relating the polarization marker iNOS to exogenous pro- and anti-inflammatory stimuli. These stimuli-to-iNOS response models were identified using linear autoregressive with exogenous input terms (ARX) equations and coupled with nonlinear elements to account for experimentally identified supra-additive and hysteretic effects. By designing transfer function models with the intent to predict cell behavior, we reproduced experimentally observed temporal iNOS dynamics induced by lipopolysaccharides (LPS) and interferon gamma (IFN-γ). Moreover, the identified model enabled the design of time-varying input trajectories to experimentally sustain the duration and magnitude of iNOS expression from both naïve and non-naïve initial states. The temporal methodology presented here will have numerous applications for regulating immune activity dynamics in chronic inflammatory diseases.

## INTRODUCTION

Healthy immune response during infection or injury is a dynamic process consisting of initial, acute pro-inflammatory activation followed by anti-inflammatory/resolving activity, which is mediated in large part by macrophages^1,2^. This temporally regulated response promotes pathogen and debris clearance followed by tissue regeneration and, ultimately, recovery of homeostasis (**Fig. 1a**)^1,2^. Dysregulation can occur in several ways. First, a strong initial pro-inflammatory response within the affected tissue can lead to systemic inflammation that positively feeds back to sustain local inflammation. Second, a compensatory anti-inflammatory response (e.g., via regulatory T cells) can lead to aberrant immunosuppression, which impairs pathogen clearance and regeneration^3,4^. Third, long-term dysregulation of immune response during chronic disease interferes with tissue regeneration and homeostasis, in turn further sustaining immune dysregulation. Indeed, chronic inflammatory dysfunction contributes to a breadth of diseases, including impaired wound healing after major trauma and multiple neurodegenerative diseases^5-7^ (**Fig. 1a**), and chronically impaired immune response can lead to worsened outcomes after new insults^8^. However, broad ablation of immune response, e.g., via corticosteroids, can equally limit successful regeneration and recovery of tissue homeostasis^6,9-12^.

**Fig. 1.**
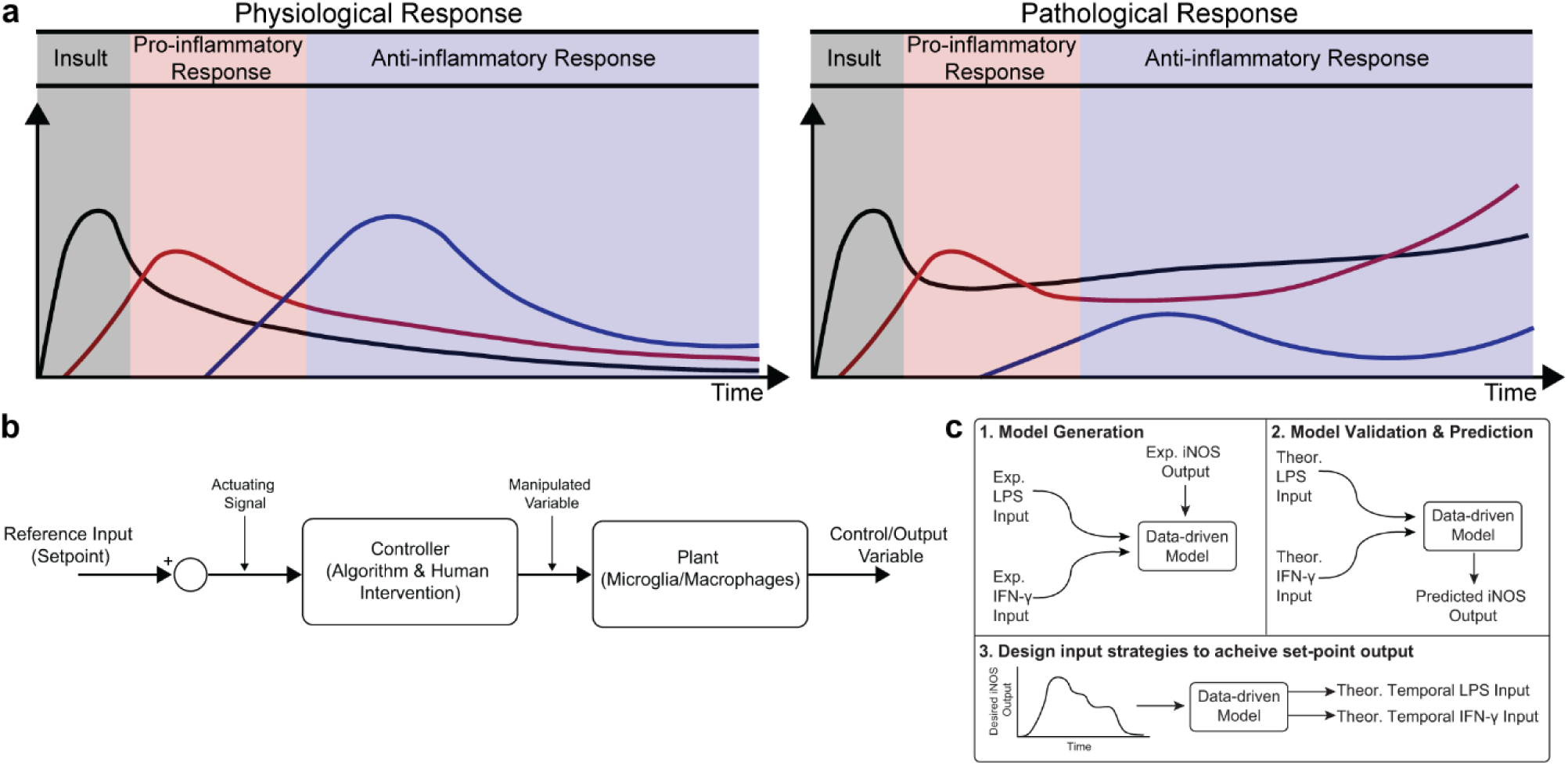
Conceptual diagram of modeling immune response in health and disease. (**a**) Immune response as dynamically regulated in health (left) and dysfunctional in chronic conditions (right). (**b**) Block diagram with macrophages as the “system” or “plant” that is being controlled. (**c**) Identification, validation and prediction of inflammatory response as a three-step process consisting of **1)** design of an engineering model structure and fit of model parameters, **2)** Comparison of predicted and experimental results, and **3)** Use of the predictive model to design input strategies to obtain a desired response.

Although the need for regulation of tissue immune response is well-recognized, identification of new strategies to intervene in tissue inflammation remains a major challenge. After trauma for example, treatment selection, dosing, and timing of administration are all crucial factors in determining patient outcome^5,13,14^. There has recently been a call for a better understanding of the complex and dynamic immune response post-injury in order to identify new strategies to regulate dynamic immune response and ultimately patient outcome^5,15,16^.

The dynamic activity of macrophages is integral to both the early (<1hr) and continued (>1 month) response to infections and injury^8,17^. Without appropriate regulation of their activity, macrophages can drive the initiation and progression of many diseases^7,8^. In particular, loss of regulation can lead to insufficient pro-inflammatory activity, leading to incomplete clearance of pathogens and/or tissue debris, impaired pro-regenerative response, chronic inflammation, and infection^6,9,18^. Recent efforts to regulate dysfunctional macrophages have focused on cell-based therapies, such as delivery of mesenchymal stem cells (MSCs) or macrophages conditioned *ex vivo* toward anti-inflammatory and pro-regenerative “M2” phenotypes. The underlying principal behind immunomodulatory cell therapies is that these cells will act as natural “controllers” of immune response through beneficial immunomodulatory signaling in the local environment^19^. However, these strategies are subject to a number of limitations. For example, MSCs are subject to variable efficacy between donors and batches^19,20^. Other approaches seek to deliver *ex vivo* modified macrophages, but both mouse and human trials have had variable success and still face many challenges^21,22^. A new approach that actively regulates resident tissue macrophages would escape many challenges faced by current cell-based therapies.

Exogenous control of macrophage activity would provide an exciting new method to modulate immune response^2,7^ that would steer the system through a desired trajectory of activity. Macrophages are an attractive target for regulating immune response because i) they are involved in diverse immune functions essential for tissue protection and repair and ii) they are highly plastic, with the ability to dynamically re-polarize for different functions based on external cues^8^. Since macrophage polarization is dynamic, a quantitative temporal model will enable design of exogenous input sequences capable of normalizing response (**Fig. 1a,b**). The pathways governing macrophage polarization in response to stimuli have been comprehensively modeled, including receptor binding kinetics, downstream kinase signaling, and gene transcription^23^. While mechanistically appealing, these models possess dozens of equations and hundreds of parameters, making it intractable to identify reliably predictive input-output relationships between exogenous stimulation and polarization in terms of these precise mechanistic models. Moreover, it has recently been argued that identification of viable strategies to intervene in immune activity will require rigorous integration of experimental data with computational modeling^24^. There is thus a need for an empirical input/output model that relates macrophage response to exogenous inputs in order to predict and control activation levels over time.

In the current study, we formulated a data-driven modeling approach, informed by an *in vitro* macrophage polarization assay and system identification theory, to identify the temporal dynamics of macrophage response to multiple exogenous pro-inflammatory stimuli. Specifically, we conditioned RAW 264.7 macrophages with M1 polarizing stimuli (LPS and IFN-γ) or an M2 polarizing stimulus (IL-4) and quantified response in terms of iNOS expression for 1-72hr post-stimulation. We then used least squares regression to fit a low-order autoregressive with exogenous terms (ARX) model together with nonlinear elements to relate iNOS response to each input (**Fig. 1c.1-2**). The identified model predicted the dynamics of polarization in subsequent experiments in response to different concentrations and temporal trajectories (simultaneous vs sequential) of each input (**Fig. 1c.3**). Finally, we used the identified model as part of an open-loop control framework to tailor input sequences to achieve desired temporal trajectories of macrophage polarization *in vitro*. To our knowledge, this is the first study to experimentally control immune cell dynamics using a predictive control framework. Given the importance of dynamic M1 and M2 polarization during tissue regeneration, the control methodology presented here defines a novel framework that will have diverse applications for treating chronic inflammatory diseases and promoting tissue regeneration.

**Fig. 2.**
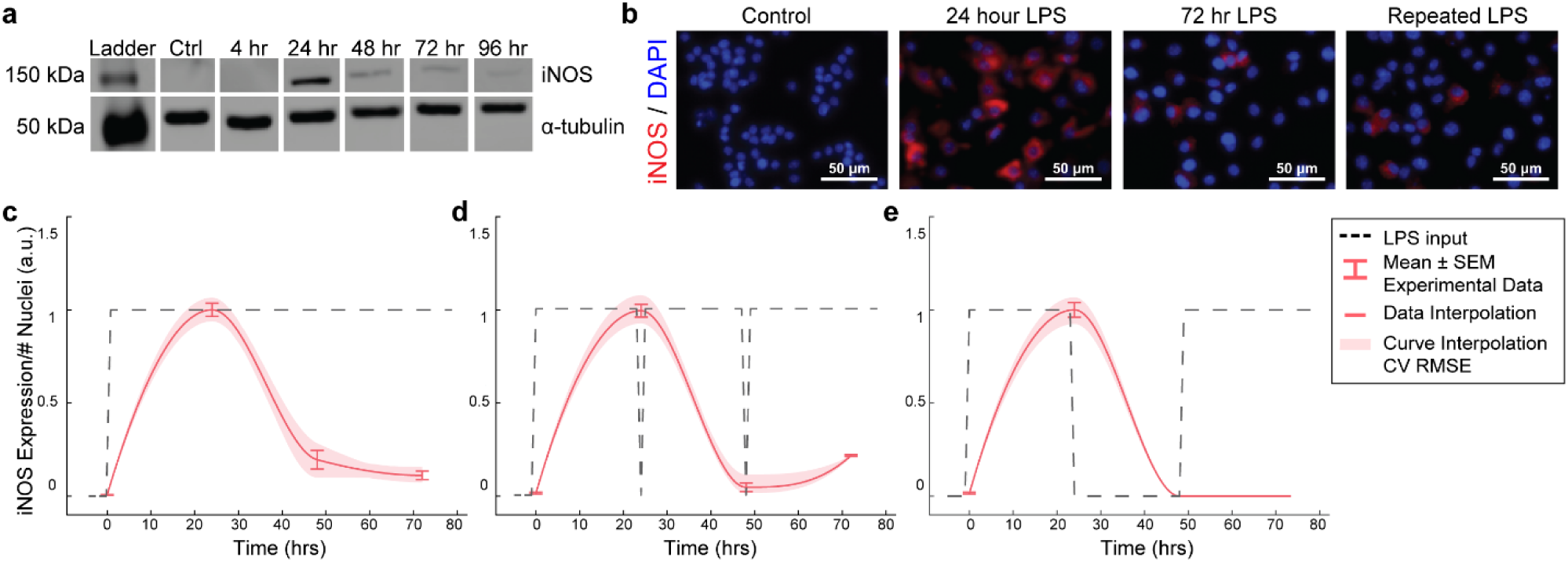
RAW264.7 macrophages transiently express iNOS in response to constant or repeated LPS stimulation. (**a**) Representative Western blot for iNOS (140 kDa) and α-tubulin (55kDa) after LPS treatment. (**b**) Representative ICC images showing iNOS response after LPS stimulation. (**c**) ICC quantification matches Western blot analysis of transient iNOS expression in response to a single administration of LPS. (**d**) Dynamics of iNOS expression are not modulated in response to multiple administrations of LPS or (**e**) after 24 hours in basal medium before LPS re-stimulation (mean±SEM, N=16 at 0, 24, 48, 72 hrs; red curves; interpolation ±RMS CV error).

**Fig. 3.**
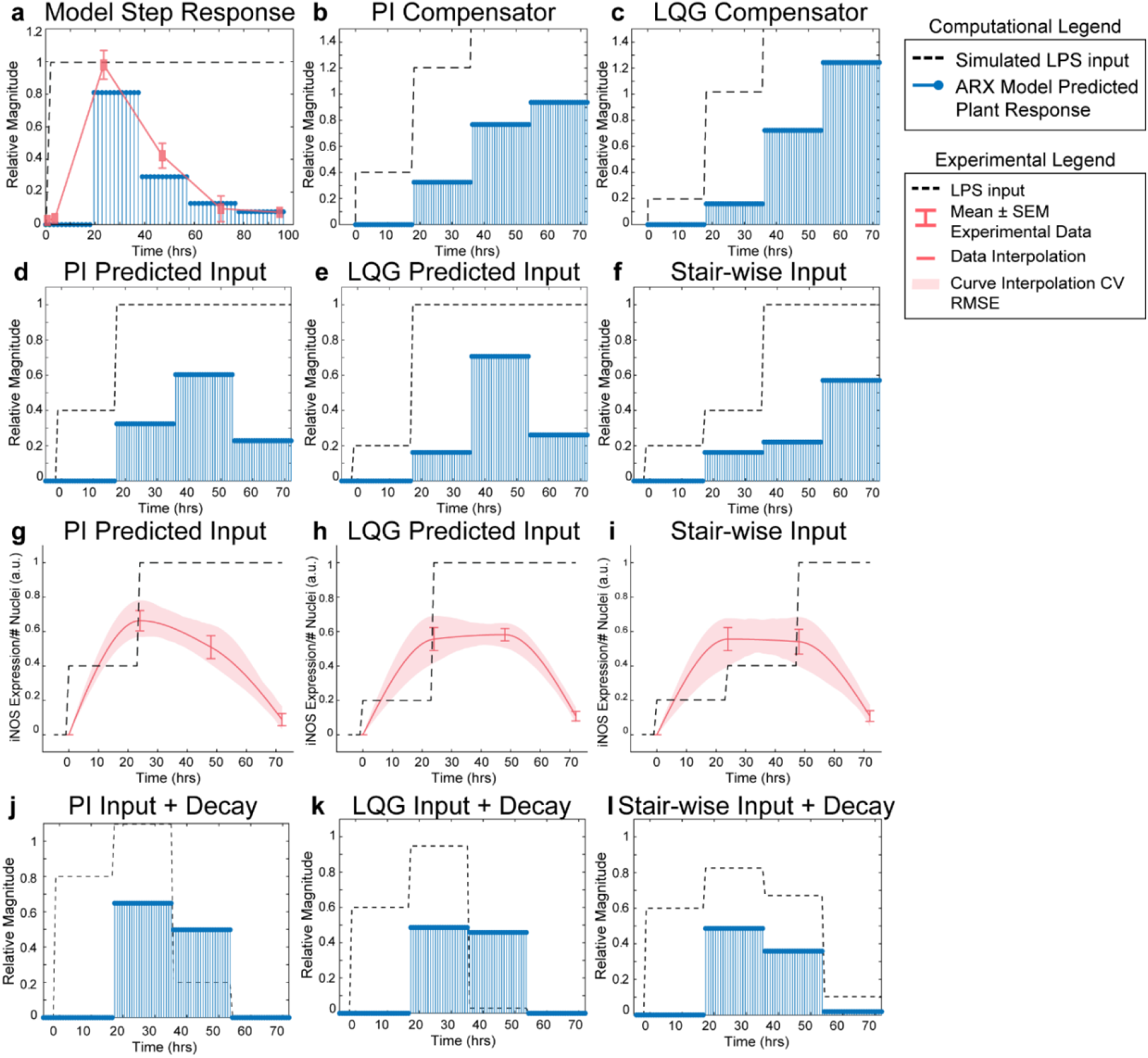
SISO LPS/iNOS ARX model, controller design, and experimental MPC testing. (**a**) Identified ARX model of macrophage iNOS response to LPS has a characteristic step response that follows the experimentally quantified trajectory. Control system design identifies input strategy (dashed line) for a step reference that elicits a gradual increase in plant response (blue stems) using a (**b**) PI or (**c**) LQG controller. Model simulations given controller defined inputs but within experimental input constraints predict sustained outputs for (**d**) PI and (**e**) LQG controllers. (**f**) A heuristically set three-step increase input strategy predicts an output that reaches a maximum at 7hr. Experimental implementation using cultured RAW 264.7 macrophages and (**g**) PI controller-, (**h**) LQG controller-, or (**i**) a heuristic combination of designed LPS input schema (dashed line) modulates temporal iNOS expression (red curves, mean±SEM, N=16; interpolated curve±RMS CV error) but does not reach the unit reference nor sustain 72 hr activity. Macrophage refractory response to repeated LPS input is captured (blue stems) by multiplying the (**j**) PI predicted, (**k**) LQG predicted, or (**l**) heuristically defined input sequences against a time-dependent exponential decay term (dashed lines).

## METHODS

### RAW 264.7 Macrophage Cell Culture and Conditioning

All studies in this work were performed using RAW 264.7 murine immortalized macrophages (ATCC TIB-71™). Macrophages were expanded, maintained, and cultured in basal macrophage medium, which is comprised of DMEM (Thermo Fisher Scientific; 12430062), 10% FBS (Thermo Fisher Scientific; 26140079), and 1% antibiotic/antimycotic (Sigma-Aldrich; A5955). Cells were cultured to 70% confluence before conditioning began. Cells were conditioned by addition of medium with lipopolysaccharide (LPS; Sigma-Aldrich; L2880), interferon gamma (IFN-γ; R&D Systems; 485-MI), or interleukin (IL)-4 (PeproTech; 214-14) as indicated. RAW 264.7 macrophages were conditioned with LPS or IFN-γ alone to quantify individual stimulus dynamic response, with LPS or IFN-γ sequentially to recover iNOS expression via orthogonal input, or with LPS or IFN-γ simultaneously to quantify supra-additivity and model predictive control strategy response. Pre-treatment, 24 hours of IL-4 prior to addition of LPS or IFN-γ, was used to induce an anti-inflammatory, non-naïve state for experiments involving hysteretic effects.

### Quantification of iNOS Expression via Immunofluorescence and Western Blot

For immunocytochemistry (ICC) experiments, macrophages were cultured in 96-well microplates. Macrophages were fixed in 4% PFA solution for 15 minutes and blocked with 5% BSA + 3% goat serum in PBS for one hour. Cells were stained with α-iNOS antibody (Cell Signaling Technology; Cat. No. 13120; 1:400) and DAPI for normalization to nuclei count. Cells were imaged at 10X magnification (Zeiss Observer Z1). Image fluorescence was thresholded and total fluorescence above the threshold was normalized to nuclei number.

For Western blot experiments, cells were cultured in 6-well plates then lysed using RIPA buffer with PMSF (Sigma-Aldrich), and cOmplete Mini (Sigma-Aldrich). Membranes were probed for α-tubulin (Sigma-Aldrich, Cat. No. T6074; 1:4000) and iNOS (1:1000). Membranes were imaged on a LiCor Odyssey CLx machine and quantified in ImageStudio Lite. iNOS band intensity was normalized to α-tubulin intensity to yield iNOS expression.

### Data Normalization and Dynamic iNOS Response Figure Generation

ICC and Western blot data were aggregated and iNOS expression for each independent experiment was normalized to the positive control with RAW 264.7 cells treated with 1μg/mL LPS for 24 hours (dynoDataLoad.m). iNOS dynamics plots were generated using the Gramm package for MATlAB^25^. Data at sampled time points (0, 24 48, and 72 hours) were expressed as mean ± SEM for separated data (N=38 for LPS single input experiments; N=8 for LPS repeated input experiments; N=8 for LPS cycled input experiments; N=32 for IFN-γ single input experiments; N=16 for IFN-γ repeated input experiments; N=16 for IFN-γ cycled input experiments). To generate interpolation curves data were smoothed using the Savitzky-Golay (sgolay) option in the curve fitting toolbox. Shaded band on curve represents root mean squared (RMS) cross validation error on smoothed data^25^.

### SISO and MISO Linear ARX Model System Identification

LPS response data were compiled into a time-domain data object with experiments for all input concentrations and unique input sequences. Dynamic models were fit (**Table S1**) to the autoregressive with exogenous inputs (ARX) model structure 

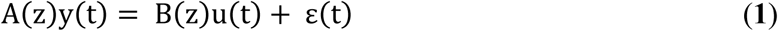

where u(t) is the LPS stimulation input, y(t) is the iNOS response, and the model coefficients consist of 

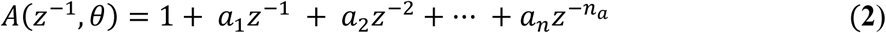

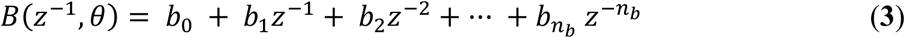

with one pole (n_a_), two zeros (n_b_), an input-output delay of 1 time step corresponding to 24hr, and zero initial conditions (System Identification toolbox, MATLAB). Parameters were estimated by solving the least squares problem 

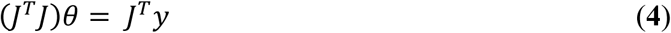

where J is the regressor matrix consisting of given inputs, *y* is the measured output, and the uniquely identified solution to the least squares parameter estimation is 

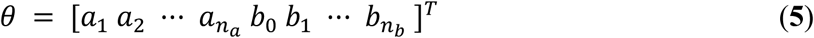

The sampling time step of the identified model was set to 24hr, which was equal to the data acquisition time step.

Realized for control design and flow diagram integration, the canonical state space equations for this ARX model are of the form **Eqs. 6 &7** with matrix coefficients listed in **Table S2**. 

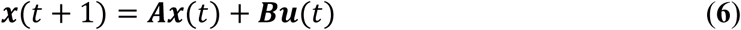

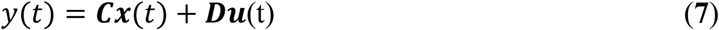

where **A** is the system matrix, **B** is the input matrix, **C** is the output matrix, **D** is the feedthrough matrix, and *t* is time. Model order was selected to minimize the small sample-size corrected Akaike’s Information Criterion (AICc)^26^ and mean squared error (**Table S3 & S4**). This process was repeated for a SISO IFN-γ model (n_a_ = 1, n_b_ = 2) and a multi-input single output (MISO) model with both LPS and IFN-γ inputs (n_a_ = 1, n_b_ = 2 for both inputs).

### LPS System Controller Design

Controller design was carried out in the Control System Designer application (MATLAB, Mathworks) to find an input strategy capable of achieving the unit step response from a step reference. Since our estimated system dynamics indicated a continuous time zero at the origin, we selected a PI controller to compensate because it adds a continuous time pole at the origin and is widely used in engineered systems^27^.A proportional-integral (PI) controller (time domain equation (**Eq. 8**) and transfer function form (**Eq. 9**), was designed with robust noise and quick response specifications (parameters given in **Table S5**). 

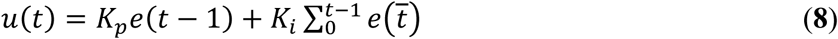

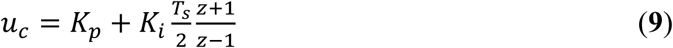

Additionally, since our system model, **Eq. 1**, enabled state estimation, we implemented a third order linear-quadratic Gaussian (LQG) controller, defined to minimize 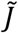 

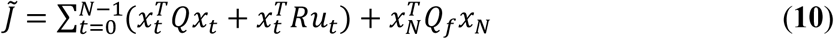

The controller was tuned to be robust to noise and assuming moderate measurement noise (zero/pole/gain parameters in **Table S6**). where N is the time horizon, t is the time step, Q is the state cost matrix, Q_f_ is the final state cost matrix, and R is the input cost matrix. Q, Q_f_, and R were defined internally by the system designer application.

### Surface Interpolation for Nonlinear Model Elements Parameterization

#### Supra-additive pro-inflammatory surface

Data matrices across concentration gradients of simultaneous LPS and IFN-γ addition were divided by the iNOS expression level given LPS only for each concentration to give the ratio by which each IFN-γ concentration amplified iNOS expression. The discrete matrix data were fit using cubic interpolation (Curve Fitting Toolbox) for each sampled time point. The resulting scaling factor, *λ*, was queried for intermediary concentrations of each input at each sampled time.

#### M2 hysteresis surface

For each LPS concentration, iNOS expression for non-M2 polarized LPS-only treated cells were divided by iNOS expression values from cells treated with an array of IL-4 concentrations for 24hr followed by 24hr of LPS. The matrix of LPS and IL-4 concentrations was interpolated using 3^rd^ order linear least squares, which provided inverse of the continuous input concentration-dependent attenuation factor γ.

### Global System Model Architecture and Formulation

For our first nested model, we used a multiple regression with interaction terms to quantify the supra-additive effect of adding both IFN-γ and LPS. Simulations were run using SISO models for single- and double-stimuli experimental results to populate a table with predicted output levels for varying magnitudes of input. The linear dual-input (both IFN-γ and LPS for all time points) model predictions were used as the regression output *y*, and the single input (either IFN-γ or LPS) SISO model predictions were given as regression inputs to fit a model of relative contributions of time and input interactions (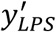 and 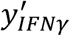). The terms that significantly predicted total iNOS output *y* were time-dependent LPS concentration, time-dependent IFN-γ concentration, and a combinatorial effect of both LPS and IFN-γ inputs (**Eq. 11**). Weighting coefficients, *c*, for each term are given in **Table S7**. 

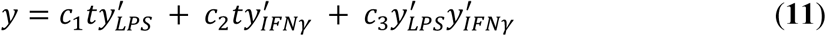

We next sought to construct a second global model structure that handles time- and concentration-dependent supra-additive interaction terms. Here, experimentally obtained iNOS expression data given varying concentrations of LPS and IFN-γ was fit to a response surface, as described above, for each time point. This surface was used to define a table as above but with time and input-dependent dual-input model output predictions. A multiple linear regression on this prediction table similarly fit coefficients for time and input interaction terms (**Eq. 11, Table S7**). We accounted for this temporally shifting interaction term by implementing the multiple linear regression model with the output from the identified SISO transfer function models *and* time as inputs and the MISO transfer function output as multiple regression model output,

### Global System Model MPC Controller Design and Prediction

The Model Predictive Control toolbox in MATLAB was used to create the controller and define manipulated input sequences for the MISO “global” model. The SISO IFN-γ and LPS transfer functions with weighting coefficients derived from the multiple regression was given as the model object, referred to as the plant (**Eq. 12, Fig. 1b**). The plant model was defined with two manipulated variable inputs, one output, a control horizon of 72 hours, and a prediction horizon of 120 hours. Manipulated variables were constrained with a minimum of 0, a maximum of 1, and unconstrained rates of change. The default state estimator (Kalman filter) settings were used for the controller predictions (MATLAB). Closed loop simulations generated the inputs, *u*, needed to obtain the set reference (unit step) over simulation time with the expected system output *y*. Plant performance was evaluated by running open-loop simulations given the predicted inputs from the closed-loop simulation. Optimal predicted input and output trajectories were validated using the mpcmove function. 

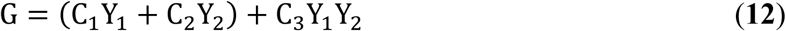

## RESULTS

### Macrophage iNOS Expression is Transient and Refractory to Repeated Stimulations

We first aimed to determine the temporal dynamics of macrophage response to single or repeated pro-inflammatory stimuli. As a model system, we used expression of the pro-inflammatory M1 marker inducible nitric oxide synthase (iNOS) by RAW 264.7 macrophages in response to the pro-inflammatory stimulus lipopolysaccharide (LPS). Using quantitative Western blot, we found that a single administration of 1μg/ml LPS, but not IL-4 (**Fig. S1**), resulted in transient iNOS dynamics with a peak in iNOS expression at 24hr followed by a decay to baseline over the following 48hr (**Fig. 2a**). Immunocytochemistry (ICC) confirmed this response (**Fig. 2b,c**) and revealed that this temporal trajectory was 1) conserved given a range of lower doses of LPS and 2) that the magnitude of the response monotonically increased with the magnitude of the stimulation (**Fig. S2**). Intriguingly, although LPS was not removed from cultures, and thus represented a persistent step-like stimulus, the dynamics of iNOS expression followed a first order decay response (**Fig. 2b,c**). In traditional engineered systems, this type of system response is usually obtained by stimulating the system with a finite impulse input^26^.

To test whether the observed decay in iNOS expression was due to LPS depletion from the culture medium, we re-administered 1μg/ml LPS every 24hr. However, iNOS expression in response to repeated stimulation was comparable to that of a single LPS stimulation (**Fig. 2d**), indicating suppression of response to subsequent stimulations. To determine if this transient response is determined by an auto-inhibitory process, we next tested the effects of cyclically re-stimulating cells with LPS following 24hr in control medium. However, cycled re-stimulation did not alter iNOS expression dynamics (**Fig. *2*e**), suggesting that the dynamics of macrophage polarization to LPS stimulation consist of an initial response that is not sustained despite either continued or repeated LPS stimulation, i.e., the system becomes refractory. This refractory behavior resembles tolerance/fatigue observed in chronic disease conditions, such as type 2 diabetes and cancer^28,29^.

### Auto-regressive Model with Exogenous Inputs Fits iNOS Dynamic Response to LPS Input

We next asked if a control systems engineering methodology could be used to design a temporal sequence of LPS stimulation that would enable us to recover and sustain iNOS expression, and, by extension, pro-inflammatory activation of RAW 264.7 cells. Control systems methodology requires a model that can be used to predict future system response given a known stimulation input. Diverse model structures are employed in engineering fields, ranging from high-ordered mechanistic models to input-output data-driven models. For this application, a mechanistic model encoding all of the genetic and protein interactions responsible for iNOS expression would suffer from reduced predictive capacity due to uncertainty in fitted parameters. Grey and black box models, which capture dominant response dynamics without specifying mechanistic details, are thus more appealing to relate iNOS dynamics to pro-inflammatory stimulation^30^. We therefore sought to identify an optimized black box single input and single output (SISO) model relating LPS input to iNOS output^30,31^. A critical tradeoff must be considered when choosing model structure: maximize flexibility to best capture system dynamics while avoiding the need to have more model parameters than can be reliably identified from the data^32^. Autoregressive models with exogenous inputs (ARX) models are frequently used for black-box system identification because they can capture underlying system dynamics in diverse applications and because parameterization using the ARX structure guarantees uniqueness of solution and identification of the global minimum of the error function^30,33-36^.

To identify the parameters of this model architecture, extensive experimental characterization of macrophage polarization dynamics with multiple input patterns and magnitudes was performed to generate a rich data set to train and identify an input/output model of iNOS expression dynamics (**Fig. 2c-e**; **Fig. S2**). We experimentally found that macrophages exhibited a monotonic LPS dose-to-iNOS response relationship within a physiologically relevant concentration range (**Fig. S2**), which is well-described using the linear ARX model structure. Above a high (1μg/ml) concentration of LPS, response tapers off, potentially due to cell death or changes in intracellular signaling activity^37^. As such, we set 1μg/ml LPS as the maximum concentration used in this study. To capture the post-LPS stimulation refractory period, we selected an ARX model order (na = 1, nb = 2) that recapitulated this refractory pattern for a step input (**Fig. 3a**). The model parameter estimates are given in **Table S1** (three free coefficients) and returned a normalized Aikike’s Information Criterion (AICc) model quality metric of 430.59 and minimized mean squared error (**Tables S3-S4**). This model outperforms the related ARMAX (autoregressive-moving average with exogenous terms) model structure with similar numbers of parameters (n_a_= 1, n_b_= 2, number of moving average coefficients n_c_= 0; AICc=501.96). By estimating this input/output model, we can achieve both high descriptive and predictive capacities.

### Model Predictive Controller Identifies LPS Stimulation Sequence to Sustain iNOS Expression

Using the identified ARX system model, we sought to tune a controller (Control System Design Toolbox, MATLAB), placed upstream of the plant (**Fig 1b**), that would predict a temporally defined LPS input strategy to overcome the persistent decay in iNOS expression. We used two controller structures to design input strategies capable of achieving sustained iNOS expression. First, since our system dynamics (**Fig. 2c**) indicated that the system model responds to the derivative of the input, we attempted to compensate for the derivative using a classical proportional-integral (PI) controller, which is commonly applied in engineering applications to minimize steady-state error^27^ (**Table S5**). Here, we used the PI controller to control LPS-induced iNOS expression to the unit reference (1 a.u. iNOS relative expression, **Methods**). The controller predicted that a stair-wise delivery of LPS (**Fig. 3b, dashed line**) would give rise to a more gradual but prolonged output response, *y*, that reached the reference by the control horizon of 72 hours (**Fig. 3b, blue stems**). Importantly, the second step in input exceeded the unit input value (corresponding *in vitro* to 1μg/ml LPS), which was the upper bound of LPS concentration used in this study. When the controller was constrained to inputs between 0 and 1 (1 μg/ml LPS) no PI controller obtained by adjusting K_p_ and K_i_ (**Methods**), was capable of defining an input sequence that both maintained a *u*≤1 μg/ml and predicted *y* to reach the reference within the control time horizon.

Due to the inability of the PI controller to identify an input sequence capable of reaching or maintaining output levels at 72 hours, we next decided to take advantage of our ARX system model to re-design the input sequence using a third order linear-quadratic Gaussian (LQG) controller (**Table S6**), which can provide improved performance over conventional PID controllers for minimizing total error^38^. This LQG controller designed a reduced magnitude for the original input followed by the unit max of LPS input (**Fig. 3c, dashed line)** to achieve 80% of reference point prior to exceeding the unit max stimulation input (**Fig. 3c, blue stems**), which the PI controller defined input could not achieve within LPS concentration constraints. However, this controller also required *u*≥1 μg/ml to reach the reference. When the input is constrained to 0≤*u*≤1 μg/ml LPS, the model simulations predicted that progressive step increases in LPS would prolong the iNOS response but not sustain it at the unit reference value (**Fig. 3d,e**). Finally, when the initial magnitudes of the LQG and PI predicted inputs were heuristically combined in a three-step increase strategy, simulations predicted a maximum response at 72 hours (**Fig. 3f**)

### Experimental Implementation of Predicted LPS Input Temporarily Sustains Macrophage iNOS Activation

Each controller above defined a temporally increasing magnitude of the stimulus *u*, or LPS concentration, where the input is increased at each time step. Experimentally, the model predicted input values represent a fraction of the normalized maximum (high) LPS concentration, 1μg/ml. For example, 0.02 is 2% of the maximum 1μg/ml, or 20ng/ml, and 0.04 is 40ng/ml as in our data used for model fitting. To test the PI controller input strategy, RAW 264.7 macrophages were treated with 40ng/ml of LPS for 24 hours, followed by 1μg/ml from hour 24 until fixation at 72 hours (**Fig. 3g, dashed line)**. Despite the controller requiring *u* of 1.2, biologically this would have led to excessive cell death, likely changing the plant response. Thus, we tested the effect of the unit max of LPS in this stair-wise input scheme. The macrophage expression of iNOS peaked at approximately 70% of normalized maximum iNOS (defined by the 24hr expression level given 1μg/mL LPS) at 24hr (**Fig. 3g, red curve**). The subsequent increase in LPS concentration delivered did not sustain this level of iNOS, which declines through the 48 and 72hr time points, but does keep levels higher (∼50% max) at 48hr than an initially high level of LPS (**Fig. 3g, red curve**).

The LQG controller predicted input, 24hr of 20ng/ml followed by 48hr at 1μg/ml LPS (**Fig. 3h, dashed line**), realized an iNOS expression level ∼60% of the reference at 24 r (**Fig. 3h, red curve**). Intriguingly, here the cells sustained this iNOS level through 48hr, but not through 72hr (**Fig. 3h, red curve**). We next heuristically combined the input strategies defined by the PI and LQG controller to test whether iNOS expression at 72hr could be sustained (**Fig. 3i, dashed line**). However, iNOS expression given this strategy reflected that of the LQG controller and did not keep activation high at 72hr (**Fig. 3i, red curve**).

The refractory, or muted, iNOS response to either high, continued or step-wise increases in LPS stimulation suggested a decaying efficacy of LPS regardless of input sequence. Indeed, when the input sequence terms were multiplied by a time-dependent exponential decay (**Fig. 3j-l, dashed lines)**, the response magnitudes reflect the experimentally obtained iNOS values (**Fig. 3j-l, blue stems**) for each input strategy. Although this single input system was unable to meet control specifications, the ability to qualitatively maintain elevated pro-inflammatory macrophage activation via our predictive control framework demonstrated an exciting feasibility of the approach that may be extendable to alternate strategies that can overcome the decaying efficacy of LPS stimulation.

### IFN-γ Stimulation Increases Reachable iNOS Trajectories and adds System Nonlinearity

We found above that single or repeated stimulation with LPS was unable to indefinitely sustain iNOS expression and that sustained expression was only partially recovered by temporally modulating the input (**Fig. 3d-i**), i.e. inflammatory activity was modulated but could not be prolonged indefinitely. In engineering systems, independent inputs increase the system rank and thereby increase state achievability. That is to say, adding a secondary stimulus that operates through separate, orthogonal means, expands the internal states and reachable output of a system^39^. Therefore, we next hypothesized that a second pro-inflammatory input would improve controllability. To test this, we used IFN-γ, which signals largely independently of LPS (**Fig. 4a**) as the second, orthogonal input because 100ng/ml IFN-γ robustly increased iNOS levels despite prior LPS input (**Fig. 4b-d**). Although we also considered TNF-α as the second pro-inflammatory stimulus, we found the iNOS response is more sensitive to IFN-γ within a physiologically relevant concentration range (**Fig. S3**). Given these findings, the use of multiple pro-inflammatory inputs is promising for toggling both the magnitude and duration of macrophage activity with greater reachability than can be achieved with a single input.

**Fig. 4.**
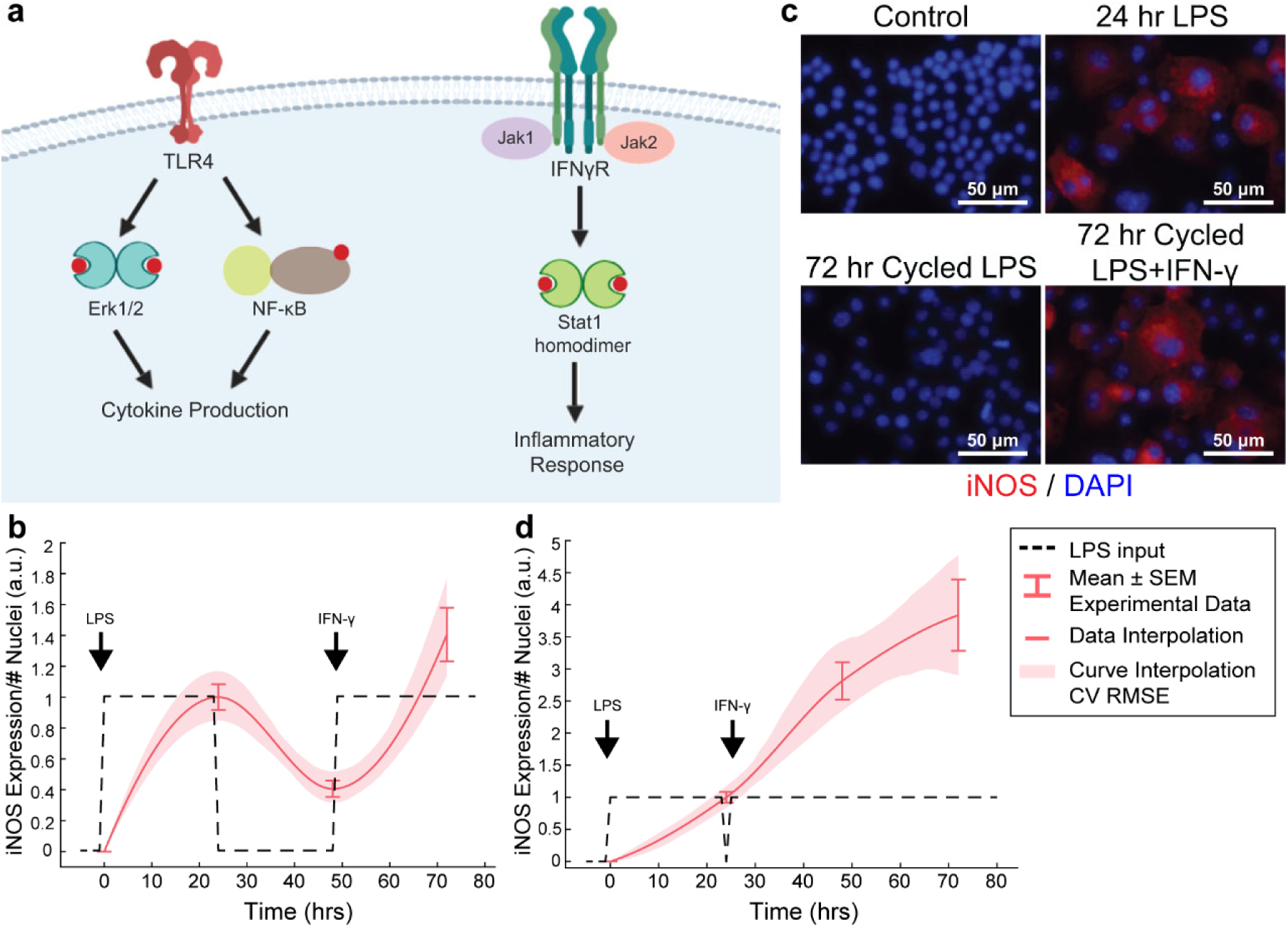
Orthogonal stimuli maintained or magnified iNOS expression. (**a**) Signaling diagram for LPS and IFN-γ (created with BioRender). (**b**) 24hrs of LPS treatment and delayed subsequent IFN-γ (dashed lines) treatment modulates iNOS expression (red curves, mean±SEM, N=16; interpolated curve±RMS CV error), even at 72hr time point. (**c**) Representative ICC images showing cycled LPS and IFN-γ (input defined in **b**) induces iNOS expression comparable to 24hr of LPS alone while cycling only LPS in that same pattern (**Fig. 2f**) does not maintain expression. (**d**) 24hrs of LPS treatment and immediately subsequent IFN-γ (dashed lines) treatment modulates iNOS expression (red curves, mean±SEM, N=16; interpolated curve±RMS CV error), even at 72hr time point.

While IFN-γ recovered iNOS expression from LPS-induced tolerance, it also introduced a non-linear element to the dynamic response – supra-additivity. ARX and transfer function models require that the output of the sum of two inputs equal the sum of the output of each input. However, IFN-γ amplifies LPS-induced iNOS expression, where expression is greater than the sum of expression from each stimulus alone, whether added concomitantly or in series. In fact, supra-additivity for simultaneous conditioning is present across all time points and for a range of LPS and IFN-γ concentrations through 72hr of conditioning (**Fig. 5a, Fig. S4**). The supra-additivity also lead to iNOS expression that was greater than the unit reference for 24 hours of LPS (**Fig. 5a**), so our predictive model needs to account for these nonlinearities to avoid overshooting or behavior that does not settled to the desired reference **Fig. 4d**.

**Fig. 5.**
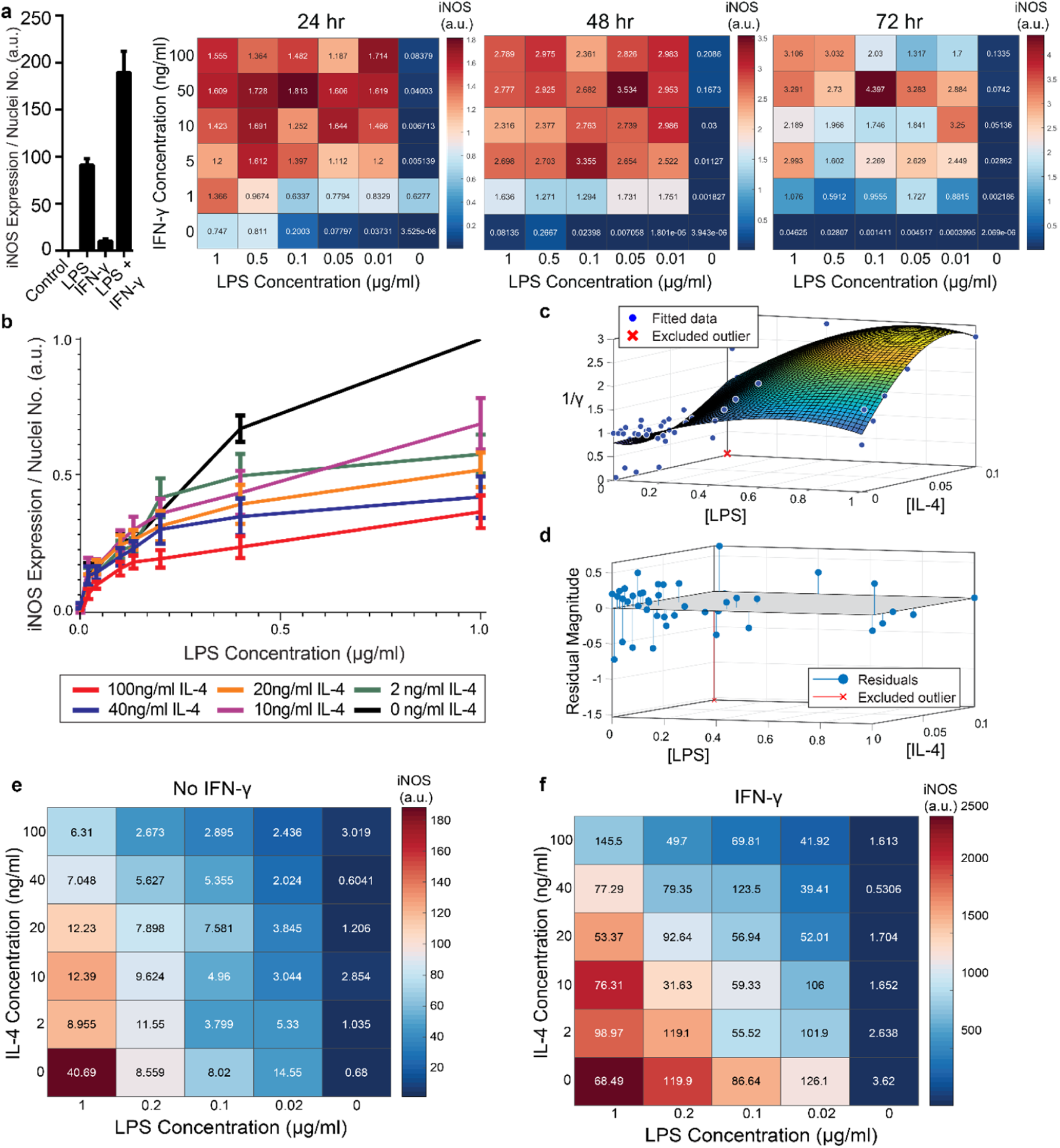
RAW 264.7 macrophages are markedly affected by activation state-dependent hysteresis, which can be overcome using multiple pro-inflammatory inputs. (**a**) LPS and IFN-γ added simultaneously cause time dependent supra-additive expression of iNOS (color and text display condition mean; N=4). (**b**) Prior treatment with IL-4 attenuates LPS induced iNOS expression (24 hr post LPS treatment) in an IL-4 concentration-dependent manner (mean±SEM, N=6). (**c**) Interpolated attenuation factor gamma surface plot and (**d**) fit error. (**e**) Pretreating macrophages with IL-4 for 24 hours prior to LPS stimulation reduced the magnitude of pro-inflammatory polarization measured by iNOS expression normalized to DAPI (color represents mean, SEM displayed numerically, N=4). (**f**) Combining 4ng/ml of IFN-γ with LPS stimulates iNOS expression, overcoming the hysteretic effect dependent on the dose of LPS (color represents mean, SEM displayed numerically, N=4).

### RAW 264.7 Macrophages Exhibit State Memory Based on Stimulation History

In disease, macrophages may exist in chronically activated or other non-naïve states, driven by local and systemic changes in signaling proteins, hormones, among other factors^7,40^. Thus, having shown our ability to model macrophage pro-inflammatory dynamics and design input trajectories for naïve macrophages, we next wanted to determine whether the macrophage response to pro-inflammatory stimulation would be affected by pre-polarizing the cells toward an anti-inflammatory state.

To model RAW 264.7 cells starting in a non-naïve state, we pre-conditioned macrophages with IL-4 for 24 hours prior to pro-inflammatory stimulation. Upon stimulation with LPS, we found that prior IL-4 conditioning attenuated expression of iNOS after 24hr of treatment with LPS, but that iNOS still responded to LPS in a concentration dependent manner (**Fig. 5b**). M2 polarization was validated by increased expression of Arg1 (**Fig. S5**). Further, an initial polarization toward a pro-inflammatory phenotype increased the magnitude of anti-inflammatory polarization that outweighed the IL-4 concentration given (**Fig. S6**), which is consistent with prior studies, including one study where AAV delivery of IFN-γ *in vivo* increased M2 gene expression, as well as M1 genes^10^. Together, these data suggest that macrophages exhibit hysteresis in their response to prior inputs, whereby prior M2 polarization attenuates future M1 response and prior M1 polarization sensitizes future M2 response. The M2 driven attenuation of M1 response reflects one aspect of how systemic immunosuppression poses a major risk to post-traumatic or surgical injury patients^41-44^.

### Modeling Multi-Input Driven Hysteresis and Supra-Additivity

Since the dynamics of iNOS expression in RAW 264.7 cells were dependent on the polarization state history (i.e., hysteresis in non-naïve cells) and demonstrated supra-additivity in response to combinations of LPS and IFN-γ, we next sought to incorporate these elements into our iNOS response model. First, to account for prior IL-4-induced hysteresis, we defined an attenuation factor to account for the relative magnitude of iNOS expression for the range of LPS and IL-4 concentrations described in **Fig. 5b** relative to expression with no exposure to IL-4. The attenuation factor, γ, is equal to 1 for non-hysteretic systems and increases with higher concentrations of IL-4 such that 1/*γ* multiplied by iNOS expression for a given LPS concentration gives the iNOS response for that LPS concentration and an IL-4 pre-treatment concentration. A response plane for γ was fitted with a 3^rd^ order by 3^rd^ order polynomial to a smoothed continuous response surface from which any attenuation due to anti-inflammatory induction is returned (**Fig. 5c,d**).

To account for supra-additive effects of multiple pro-inflammatory inputs, as done for the hysteretic surface, we populated time-dependent interaction term (λ) surface curves for the defined ranges of co-addition of LPS and IFN-γ. Excitingly, the supra-additivity of IFN-γ with LPS demonstrated the ability to recover the attenuation effect induced by IL-4. Indeed, greater iNOS expression was observed across lower LPS concentrations and higher IL-4 concentrations when IFN-γ co-stimulation was used compared with LPS stimulation alone (**Fig. 5e,f**, note that the scale of response is an order of magnitude greater in panel **f**). This interaction effect motivates the need for a system plant model that processes both M2 and M1 inputs.

The global plant model was constructed and is described schematically in **Fig. 6**. The system receives the concentration of LPS (u_1_) and IFN-γ (u_2_) which are passed into their respective identified ARX models, the supra-additivity of LPS and IFN-γ was accounted for using λ, the pro-inflammatory contributions are summed and applied as inputs to the hysteresis term γ, Finally, the output is the predicted iNOS output (*ŷ*) as a function of time *t* (**Fig. 6**).

**Fig. 6.**
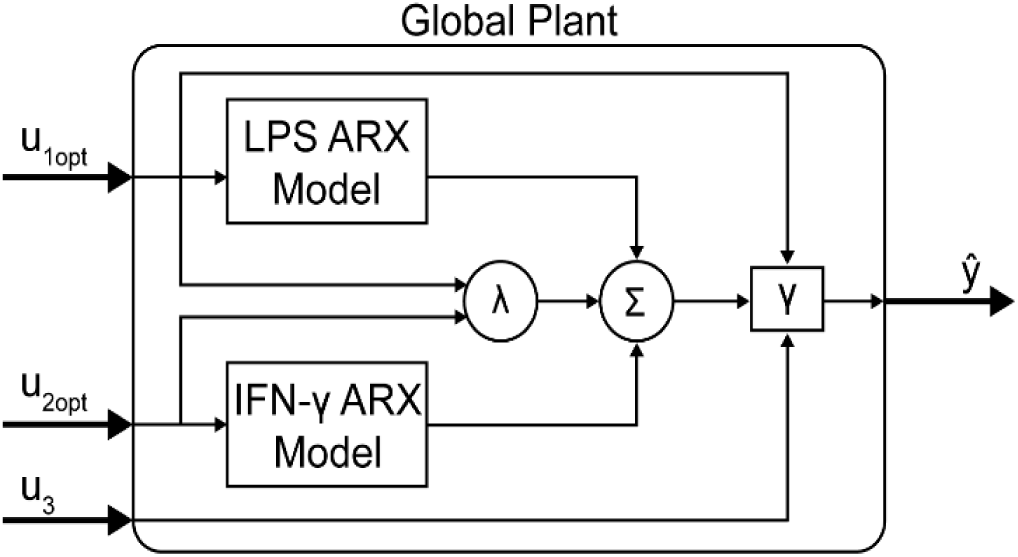
Global plant consisting of linear and non-linear model elements. Detailed diagram of multiple input plant model implemented in control system (**Fig. 1b**). System predicted inputs *u*_*1*_ (LPS) and *u*_*2*_ (IFN-γ) are fed into respective identified SISO ARX models and supra-additive interaction term *λ* elements. Terms multiplied by weighting coefficients *c* (defined by multiple regression estimation) prior to summation (*Σ*) and hysteresis-dependent attenuation (*γ*). Note that *u*_*3*_ accounts for IL-4 attenuation via *γ*.

### Design of LPS and IFN-γ Temporal Input Trajectories with Global Plant Model Achieves Sustained iNOS Expression

Transfer functions were linearly combined with regression coefficients for supra-additivity (λ) and hysteresis (γ) acting as pre-processing filters, i.e. the terms were multiplied with each model’s output, then added. The global regression of the function has the final form in **Eq. 11** (R^2^ = 0.748; p-value (vs. constant model) = 1.34e-38). Simultaneous administration of unit, high, inputs *in vitro* vastly overshot the unit value of iNOS and did not settle over the course of the experiment (**Fig. 7a**), demonstrating that it is possible to obtain sustained iNOS response, but that more carefully crafted input sequences are needed to obtain constant, sustained expression of iNOS. We therefore next used the global model (**Fig. 6**) together with an MPC controller to design input trajectories for LPS (u_1_) and IFN-γ (u_2_) needed to obtain sustained constant iNOS expression over a 72hr control horizon (**Fig. 7b**). Using these trajectories, the simulated plant reached the reference value by 24hr with a minor overshoot that settled by 72hr (**Fig. 7c**). Including hysteresis in the plant controller estimation increases the predicted inputs magnitude needed to obtain the unit step reference (**Fig. 7d**). Given the input sequence defined in (**Fig. 7d**), a hysteretic system was predicted to respond with relatively small overshoot and error (**Fig. 7e, red curve**). Importantly, the model captures the large overshoot that would be expected from administering elevated input levels to a non-hysteretic system (**Fig. 7e, blue curve**).

**Fig. 7.**
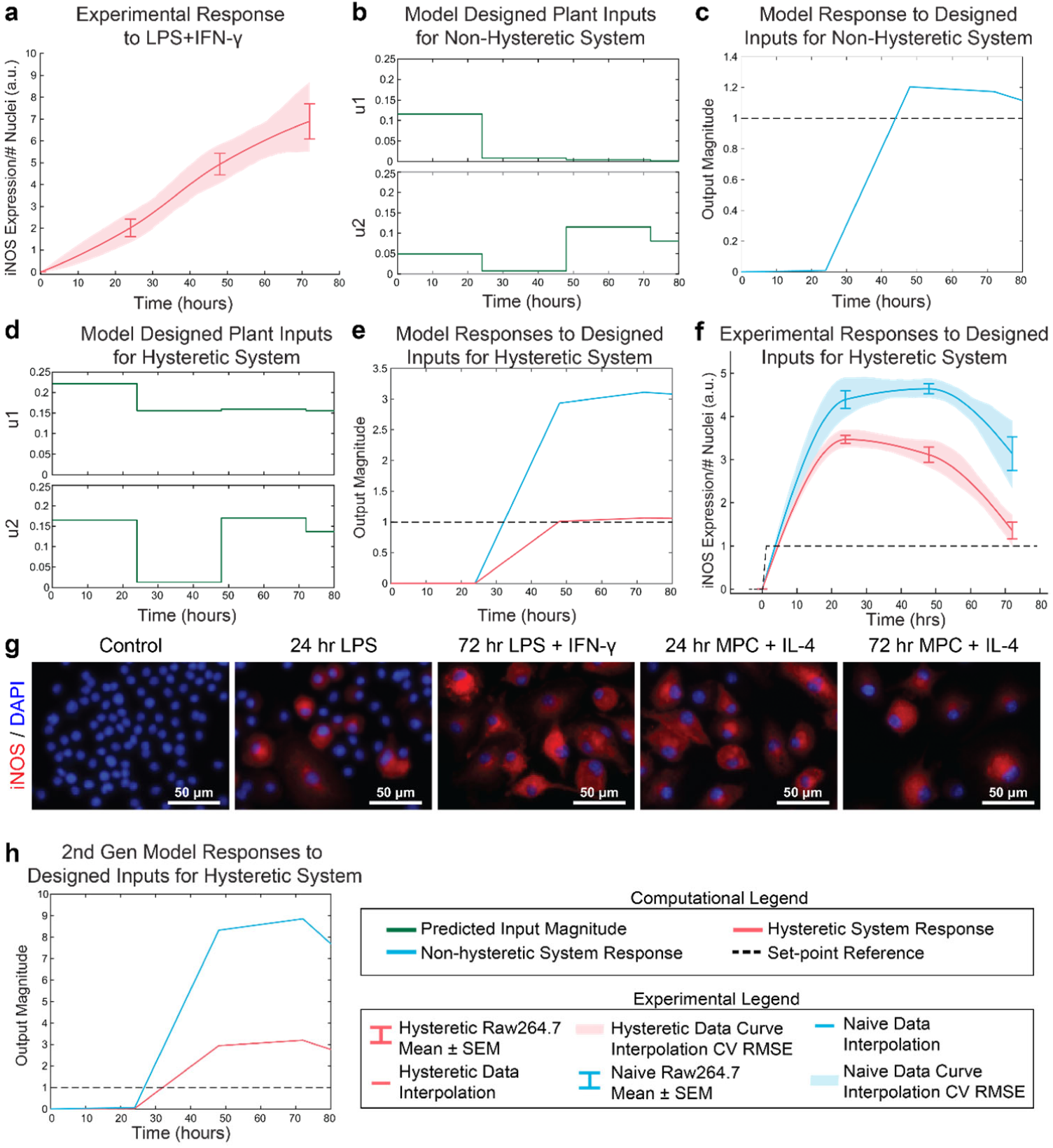
Open-loop control of pro-inflammatory macrophage activity is experimentally achieved using a nested multiple regression. (**a**) RAW 264.7 macrophage temporal response to 1μg/ml LPS and 100ng/ml IFN-γ. (**b**) Model designed inputs *u*_*1*_ and *u*_*1*_ using hysteresis-free model, which reflects cells beginning in a naïve state. (**c**) Hysteresis-free model response to inputs defined in **b**). (**d**) Model designed inputs *u*_*1*_ and *u*_*2*_ using first generation model accounting for hysteresis, which reflects cells starting from a non-naïve 24hr IL-4 primed state. (**e**) Hysteretic model (red) and non-histeretic model (blue) responses to inputs defined in **d**). (**f**) Experimental delivery of designed inputs in **d**) reflects predicted control output (**e**) for both hysteretic IL-4 primed (red curve, mean±SEM, N=16; interpolated curve±RMS CV error) and non-hysteretic (blue curve, mean±SEM, N=16; interpolated curve±RMS CV error) RAW 264.7 macrophage cultures. (**g**) Representative images of iNOS staining in model predictive control experiments using the inputs in **d**). (**h**) Simulation of updated 2^nd^ generation model with dynamic supra-additivity term in response to designed inputs (**d**) captures experimental RAW 264.7 iNOS expression for both hysteretic (red curve) and non-hysteretic (blue curve) systems.

Next, the relative input magnitudes defined for a hysteretic plant (**Fig. 7d**) were translated to concentrations of LPS and IFN-γ, which were administered as temporally defined to RAW 264.7 macrophages in culture. The macrophage iNOS expression trajectories reflected the model predicted response for both hysteretic, i.e. pretreatment with IL-4 (**Fig. 7f, red curve & 7g**) and non-hysteretic (**Fig. 7f, blue curve**) cell conditions. Since this initial model only accounted for a static supra-additivity term, we next updated it to incorporate a dynamic supra-additivity λ term that updated with time based on our response data in **Fig. 5a**. The updated model was simulated with inputs used experimentally (**Fig. 7f**) and defined by the original model (**Fig. 7d**). This 2^nd^ generation model improved the predictive performance with results that recapitulated the minor overshoot seen in the hysteretic system (**Fig. 7h**).

In total, these experimental findings show that our global plant model predicts the dynamics macrophage pro-inflammatory response, including transient response to LPS, supra-additivity, and hysteresis. Moreover, we showed that this model could be used to define dual stimulation strategies that could prolong RAW 264.7 cell polarization as quantified by iNOS.

## DISCUSSION

In this work, we developed a novel paradigm for engineering immune activity by defining predictive data-driven models of macrophage polarization and using them to define the dynamic delivery of pro-inflammatory factors to control the duration and magnitude of macrophage polarization. Rather than identifying detailed, highly parameterized mechanistic models, we applied a control theory framework to globally describe the pro-inflammatory activity of macrophages over time. Specifically, using expression of canonical pro-inflammatory (M1) marker iNOS as an output, we defined a black-box transfer function to capture the dynamic response of macrophages given a temporal sequence of applied LPS and IFN-γ as system inputs. Our overall modeling framework coupled linear ARX models, which are uniquely identifiable, with nonlinear elements that accounted for state-history dependent hysteresis and supra-additivity from multiple pro-inflammatory stimuli. Our global plant model structure not only predicted responses to different input sequences but enabled design of new stimulation sequences that yielded a desired temporal iNOS response without a refractory response (**Fig. 7**).

Immune dysregulation plays a central role in diverse diseases. Dysregulated activity of macrophages in particular can both hinder tissue repair and promote disease pathogenesis. However, macrophage functional diversity and broad distribution throughout the body also makes them excellent targets for modulating immune function to treat an array of diseases^23^. Yet the vast majority of new immunomodulatory strategies, including inflammatory inhibitors and cell-based therapies, do not explicitly account for the temporal evolution of macrophage response needed to resolve the response to injury.

The importance of a temporally dynamic immune response has been highlighted by recent findings that long term resolution of inflammation depends on a sufficiently pro-inflammatory initial response followed by anti-inflammatory and resolving activity^16^. Early pro-inflammatory macrophage response enables clearance of pathogens and damaged cells and subsequently triggers the anti-inflammatory and pro-regenerative response^21,22,45^. Thus, in the current study, we sought to model and control macrophage pro-inflammatory activity, measured by iNOS expression. Using an ARX model structure, which is widely used for black-box system identification in engineering^31^ and biological systems^30,33-36^, we identified computational models able to predict and control temporal iNOS expression. This black-box approach enabled us to fit three parameters to model the dynamic LPS response and three more to fit the IFN-γ response, in contrast to dozens required in mechanistic differential equation models of macrophage polarization^23^.

Interestingly, when implementing model-predicted LPS input sequences, we observed that the time-dependent decay in the efficacy of LPS persisted. In fact, when the designed input magnitude was multiplied against a time-dependent decay term (**Fig. 3j, l; dashed lines**), we were able to simulate the observed experimental response. This finding is consistent with macrophage auto-regulatory processes that prevent runaway inflammatory activity to LPS ^37^

The current work has some limitations that invite the need for future studies. First, we used murine RAW 264.7 immortalized macrophages, considered one of the best macrophage cell lines, for development of the methodology in this study, due to their high reproducibility between labs and studies^46,47^, but future work is needed to validate and tune the models for primary isolated macrophages. Further, to extend the utility of the model for disease therapeutics, it will be necessary to identify similarities and differences between primary macrophages collected from wild type mice and mouse models of chronic inflammatory disease. For example, macrophages are known to exhibit distinct inflammatory profiles from diabetic patients than from healthy individuals^48^, which will be reflected in the identified model parameters. Additionally, the methodology developed here lays a foundation for dynamic control of macrophage activation using a single polarization marker, but a wider panel of pro- and anti-inflammatory markers are needed to fully delineate macrophage activation state and effector function.

Together, our dynamic experimental and computational approach establishes a new way of conceptualizing and modulating macrophage activity by using a temporal sequence of input stimuli to shape the trajectory of inflammatory response. We experimentally validated the computational model predictions, extending previous theoretical work in model predictive control for patient-specific therapeutics^49^. We envision this framework having broad-reaching applications both *in vitro* an *in vivo*. Moreover, our ability to modulate macrophage activity suggests that design of temporally varying inputs has therapeutic potential for broad chronic inflammatory disorders.

## ACKNOWLEDGEMENTS

This work was supported by startup funds from the George W. Woodruff School of Mechanical Engineering at the Georgia Institute of Technology. L.D.W. was supported in part by the National Institutes of Health Cell and Tissue Engineering Biotechnology Training Grant (T32-GM008433).

## AUTHOR CONTRIBUTIONS

L.D.W and L.B.W designed the study. L.D.W, J.E.F, and A.W conducted experiments and data analysis. L.D.W conducted computational predictive modeling. J.U. conceptualized physical interpretation of control laws. All authors contributed to article preparation. L.B.W supervised the project.

## DATA AVAILABILITY

All data are available upon request.

