## Supplementary Figures and Tables for "Experimental control of macrophage pro-inflammatory dynamics using predictive models"

**Weinstock *et al.***

### SUPPLEMENTARY FIGURES AND TABLES

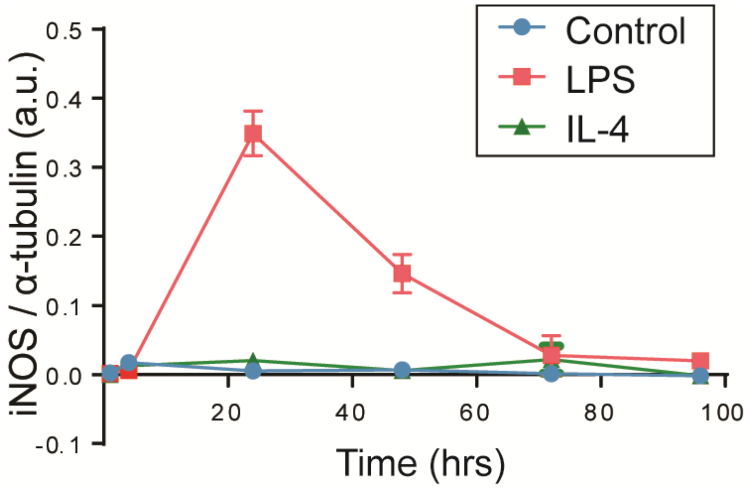


**Supplementary Figure 1.** Western blot quantification of in vitro Raw264.7 macrophage iNOS protein expression after treatment with LPS, IL-4, or control media shows iNOS peaks at 24 hrs of LPS treatment but is not expressed in IL-4 conditions (n=2; mean±min/max range).

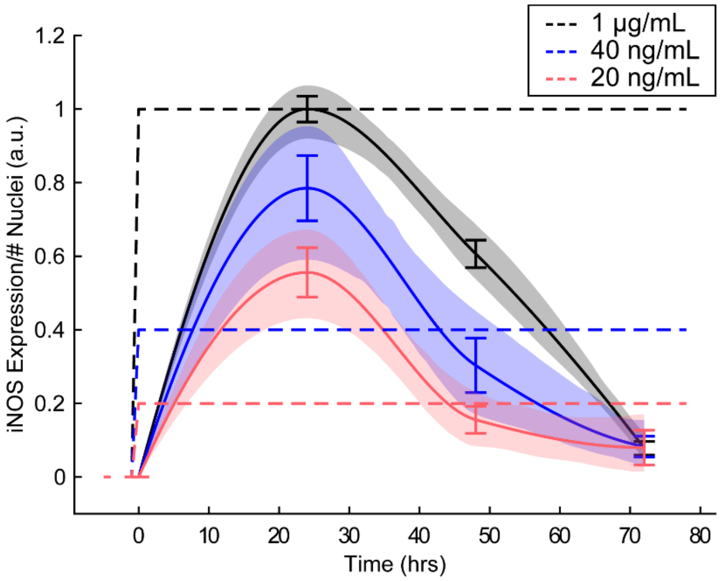


**Supplementary Figure 2.** Temporal dynamics of iNOS expression are conserved across LPS concentrations and magnitude of response is monotonic. Dashed lines represent relative LPS input. iNOS expression are mean±SEM, N=24. Solid curves were interpolated with shaded region showing RMS CV error.

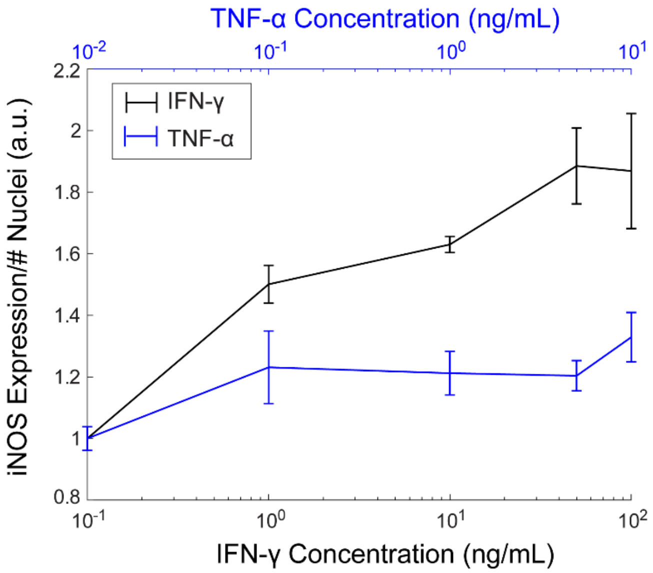


**Supplementary Figure 3**. Choice of orthogonal input.

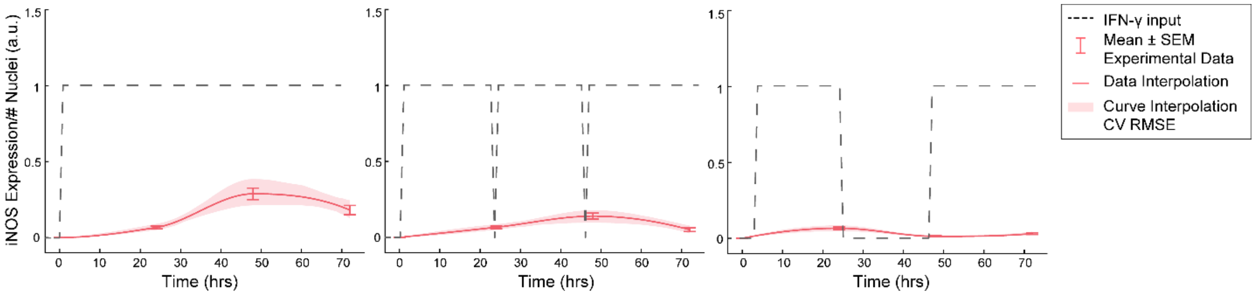


**Supplementary Figure 4.** RAW 264.7 macrophage temporally dynamic response to 100ng/ml IFN-γ alone is distinct from the LPS response but is also not sustained.

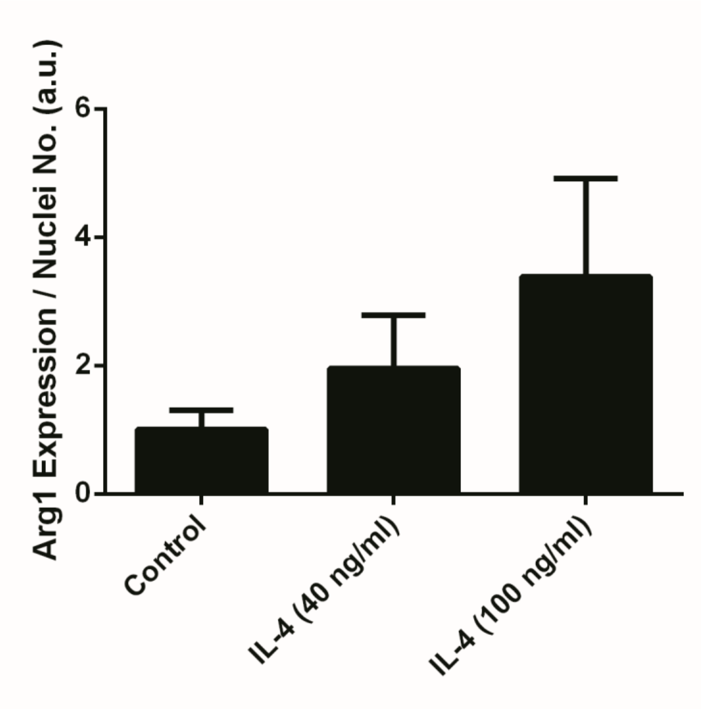

**Supplementary Figure 5.** Arg1 expression for hysteresis M2 polarization validation.

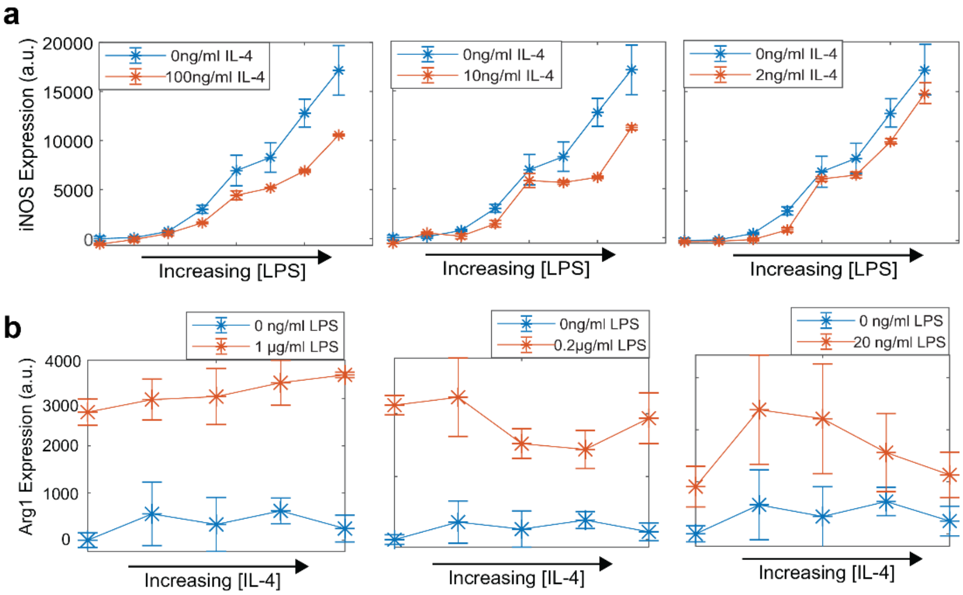


**Supplementary Figure 6. Prior M1 state polarization increases, rather than decreases, subsequent M2 polarization.** Macrophages were treated with a range of LPS concentration for 24 hrs to induce initial M1 polarization. Subsequently, a range of IL-4 doses were added for all LPS concentrations. (**a**) Increasing IL-4 concentrations attenuated iNOS expression for high LPS concentrations. (**b**) Addition of LPS stimulates Arg1 expression compared to conditioning with 24 hrs of IL-4 alone, showing a primed polarization toward M2 given prior M1 activation (**B**).

**Table S1. LPS ARX polynomials**

| | | $z^0$ | $z^{-1}$ | $z^{-2}$ |
|---|---|---|---|---|
| **LPS** | A | 1 | -0.3163 | --- |
| | B | 0 | 0.81 | -0.7727 |
| **IFN-γ** | A | 1 | -0.3849 | --- |
| | B | 0 | 0.0634 | 0.0566 |
| **LPS + IFN-γ** | A | 1 | -0.76 | --- |
| | B1 | 0 | 0 | 1.252 |
| | B2 | 0 | 2.019 | 0 |

**Table S2. LPS transfer function**

| | | **LPS Model** | **IFN-γ Model** | **LPS + IFN-γ Model** |
|---|---|---|---|---|
| **A** | $A_{11}$ | 0.3163 | 0.3849 | 0 |
| | $A_{12}$ | 0 | 0 | 0 |
| | $A_{21}$ | 0.5 | 0.5 | 1 |
| | $A_{22}$ | 0 | 0 | 0.76 |
| **B** | $B_{11}$ | 2 | 0.5 | 0.6262 |
| | $B_{12}$ | --- | --- | 0 |
| | $B_{21}$ | 0 | 0 | 0 |
| | $B_{22}$ | --- | --- | 1.009 |

| | | | | |
|---|---|---|---|---|
| **C** | $C_1$ | 0.405 | 0.1268 | 0 |
| | $C_2$ | -0.7727 | 0.2263 | 2 |
| **D** | $D_1$ | 0 | 0 | 0 |
| | $D_2$ | --- | --- | 0 |

**Table S3. LPS ARX model AICc for parameter number, $n_a$ and $n_b$, ranging from 1-4.**

| | | $n_a$ | | | |
|---|---|---|---|---|---|
| | AICc | 1 | 2 | 3 | 4 |
| $n_b$ | 1 | 331.62 | 430.59 | 548.77 | 707.25 |
| | 2 | 425.95 | 383.23 | 561.34 | 697.86 |
| | 3 | 550.49 | 562.75 | 574.56 | 711.95 |
| | 4 | 640.84 | 683.44 | 697.82 | 1789.39 |

**Table S4. LPS ARX model MSE parameter number, $n_a$ and $n_b$ ranging from 1-4.**

| | | $n_a$ | | | |
|---|---|---|---|---|---|
| | MSE | 1 | 2 | 3 | 4 |
| $n_b$ | 1 | 0.10 | 0.01 | 0.12 | 0.22 |
| | 2 | 0.04 | 0.01 | 0.12 | 0.21 |
| | 3 | 0.14 | 0.13 | 0.13 | 0.21 |
| | 4 | 0.17 | 0.20 | 0.20 | 0.21 |

**Table S5. PI controller parameters**

| | **Value** |
|---|---|
| $K_p$ | 0.401 |
| $K_i$ | 0.0334 |
| $T_s$ | 24 |

**Table S6. LQG controller design**

| | **Value** |
|---|---|
| **Zeros** | -2.631; 0.4089 |
| **Poles** | 1.0; 0.9539 |
| **Gain** | 0.1966 |

**Table S7. Multiple regression 1 interaction terms, coefficients, and p-values.**

| Term | Estimate | p-value |
|---|---|---|
| **Time : LPS-induced iNOS** | 0.1696 | 1.299e-08 |
| **Time : IFN-γ-induced iNOS** | 0.3458 | 6.485e-07 |
| **LPS-induced iNOS : IFN-γ-induced iNOS** | 69.738 | 7.366e-11 |
